# Functional Phenotyping of MMV Pandemic Response Box Identifies Stage-specific inhibitors Against Blood stage *Plasmodium*

**DOI:** 10.1101/2024.10.08.617232

**Authors:** TP Akhila, KM Darsana, Rajesh Chandramohanadas

**Affiliations:** Laboratory of Red Cell Diseases, Division of Pathogen Biology, BRIC-Rajiv Gandhi Centre for Biotechnology, Thiruvananthapuram; Manipal Academy of Higher Education, Manipal, Karnataka

**Author notes:** **Correspondence to:** Dr. Rajesh Chandramohanadas Principal Investigator, Laboratory of Red Cell Diseases, Division of Pathogen Biology, BRIC-Rajiv Gandhi Centre for Biotechnology, Thiruvananthapuram.

**Keywords:** *Plasmodium falciparum*, Pandemic Response Box, Small Molecule Inhibitors, Phenotype-based screening, drug repurposing

## Abstract

Wide-spread resistance to clinically used antimalarials necessitates the prioritization of novel scaffolds with alternate mechanisms, as possible partner drugs to artemisinin. We utilized the Pandemic Response Box chemical library of the Medicines for Malaria Venture launched in 2019 to identify inhibitors with stage-specific potency and phenotypic signatures against *P. falciparum* towards exploring the possibility of drug repurposing. From this screening, we initially identified 60 molecules active against both drug sensitive (3D7) and chloroquine resistant (Dd2) strains of *P. falciparum*. Further, 28 active compounds active below 3µM were prioritized several of which specifically impaired stage-transitions of ring (MMV001014), trophozoite (MMV1593540 and MMV1634402) and schizonts (MMV1580844, MMV1580496, MMV1580173 and MMV1580483) confirmed through microscopic phenotypes and flow cytometry. The ring stage inhibitor, MMV001014, was irreversible, led to no recrudescence and showed antagonistic effects with artemisinin indicative of overlapping mechanism. Both the trophozoite inhibitors exhibited nanomolar IC_50_ with non-compromised digestive vacuole. MMV1593540 was partially additive with artemisinin while antagonistic with chloroquine. Two among the schizont stage inhibitors (MMV1580844 and MMV1580496) appeared to operate through a mechanism driven by the generation of reactive oxygen species and all of them with molecule-specific effect on infected red blood cell (RBC) membrane integrity confirmed through confocal microscopy. Taken together, these results highlight interesting starting points derived from MMV’s Pandemic Response Box for repurposing to combat Malaria that continues to morbidly affect the developing world.

**Importance:** *Malaria* caused by infectious parasites belonging to the *Plasmodium* family continues to morbidly affect the marginalized populations. The situation is further complicated by lack of mass vaccination, drug resistance, and emergence of new parasitic forms. To alleviate the threat of drug resistance, it is important to identify new drugs acting through mechanisms distinct from the existing ones such as artemisinin. This work describes the screening of a chemical compound library against blood stage development of malaria parasites and prioritization of molecules that can inhibit parasite development in a stage-specific manner. Several of these compounds demonstrate nanomolar potency against sensitive and resistant forms of the parasites acting through distinctive mechanisms. Exploring the modes of action of these molecules will facilitate their optimization and possible clinical applications against the deadly diseases, *Malaria*.

## INTRODUCTION

Malaria caused by protozoan parasites of the species *Plasmodium,* among which *Plasmodium (P.) falciparum* is the most lethal form, leads to severe disease manifestations and economic burden to the developing world. In 2022, nearly 250 million cases were reported in 85 malaria-endemic countries with an estimated 608,000 deaths, mostly among pregnant women and children under the age of five (1). While WHO sub-Saharan Africa remains the most impacted, India and Indonesia account for 94% of WHO South East Asia’s malaria deaths (1). Since the implementation of mass vaccination being at the early stages, chemotherapy remains the primary intervention strategy in allaying the disease burden (2), primarily through *artemisinin* and its combinations (3). However, wide-spread multi-drug resistance underlines the need for new medicines with novel modes of action to combat intra-erythrocytic development of *Plasmodium* (4).

Morphological characteristics during erythrocytic development of *Plasmodium*-infected erythrocytes can be efficiently captured through phenotype-based screens *in vitro* (5), which allows for early prioritization of small molecule inhibitors. Compared to target-based screens, phenotypic screening does not require prior knowledge of the cellular target(s) and is able to identify drugs acting with multiple targets (6–8). Such whole organism-based readouts can often provide early cues on the likely mode of action, susceptibility and kinetics using which the pharmacophore could be optimized, limiting the risk of failure in the downstream translational pipeline (9–11). Literature indicates that more number of first-in-class antimalarials were discovered through phenotypic screening (6) rather than targeted approaches.

In this work, we resolved the stage and phenotype-specific inhibitory activity of compounds from the Medicines for Malaria Venture (MMV) Pandemic Response Box (PRB) against the intra-erythrocytic development of *P. falciparum* (12). PRB was launched through a Public-Private Partnership in 2019 by MMV as a way to expedite development of new drugs against infective and neglected diseases (12), by way of re-purposing. This library contains 400 structurally diverse compounds, among which 201 possess antibacterial activity, 153 antiviral, and 46 antifungals. The PRB library also contains several reference compounds with proven activity against one or the other infectious agents. In prior work, PRB has been utilized to identify inhibitors against clinically relevant bacterial, fungal, and protozoan pathogens, such as *Klebsiella pneumonia, Pseudomonas aeruginosa*, and *Schistosoma mansoni* (13–15). Compounds showing activity against multiple stages of *Plasmodium* (asexual, liver and gametocyte stages) have been identified from the PRB through *in vitro* screening approaches (16). Furthermore, novel inhibitors of *Pf*ATP4, a Na^+^ efflux pump of the *Plasmodium* parasites were reported through a structure-based virtual screening method (17).

To expand our current understanding on PRB molecules with potency against blood-stage *Plasmodium* and to capture the phenotypes indicative of possible modes of action, we performed a systematic cross-screening against drug-sensitive (3D7) and CQ-resistant (Dd2) parasites. From this, 28 compounds with dose-dependent inhibitory activity were identified. Through life-stage specific inhibition assays, we identified MMV001014 as a potent ring stage inhibitor, which demonstrated irreversible growth perturbation activity with nanomolar efficacy. Complete parasite clearance with no recrudescence was observed even after 14 cycles indicating the suitability of MMV001014 for possible clinical exploitation. Similarly, our screens returned MMV1593540 and MMV1634402 as inhibitors of trophozoite development and four molecules (MMV1580844, MMV1580496, MMV1580173 and MMV1580483) inhibiting the schizonts. These stage-specific molecules were further examined for their phenotypic uniqueness, ability to induce ROS generation, combinatorial activity with *chloroquine* and *artemesinin* as well as their ability to kill *artemisinin*-resistant (*Pf*R539T) parasites. Understanding the molecular basis of these growth perturbations and underpinning biology will unravel novel scenarios for antimalarial drug discovery.

## MATERIALS AND METHODS

### Ethics statement

All experimental procedures were conducted in accordance with the approved biosafety guidelines of the Institutional Bio-Safety Committee (IBSC), BRIC-Rajiv Gandhi Centre for Biotechnology (RGCB). Blood collection protocol from healthy volunteers for culturing *P. falciparum* was approved by the Institutional Human Ethics Committee.

### Blood collection and storage

Blood collected in EDTA tubes (BD Vacutainer) was washed thrice in incomplete media (RPMI with HEPES (Gibco) along with Hypoxanthine-Sodium hydroxide-Sodium bicarbonate and Gentamicin (Sigma-Aldrich)) to remove the buffy coat. Washed RBCs were stored at 4°C in incomplete media at 50% haematocrit for up to 3 weeks for parasite culturing.

### Preparation of working concentrations of PRB compounds

The PRB library was received from MMV (Geneva, Switzerland) in lyophilized form, which were reconstituted in 100µl DMSO to a stock concentration of 1mM and stored at −80°C until further use. Aliquots were maintained separately to avoid frequent freeze and thaw to not compromise the compound integrity. Working stocks of the compounds were appropriately made by further dilutions in DMSO.

### Parasite culture, synchronization and maintenance

We used 3D7 and Dd2 strains of *P. falciparum* (MR4) unless otherwise mentioned. Parasites were cultured in complete Medium (RPMI/HEPES with Hypoxanthine-Sodium hydroxide - Sodium bicarbonate, Gentamicin (Sigma-Aldrich), supplemented with Albumax (Gibco)) in a CO_2_ incubator as described in prior work (18). Synchronization was performed either at the ring stage using 5% sorbitol (Sigma-Aldrich) or at the schizont stage through magnetic selection (MACS; Miltenyi Biotec) (18, 19). Parasitaemia was scored using flow cytometry (FACS Aria 3, BD Biosciences, USA) on glutaraldehyde-fixed samples, which were permeabilized and stained with Hoechst 33342. In parallel, Giemsa-stained smears were examined through light microscopy for phenotypic characterization. All experiments were performed at least three times with three technical replicates.

### Profiling of PRB library to determine active compounds against *Plasmodium*

Sorbitol synchronized 3D7 parasites were seeded at 2% parasitemia at 2% hematocrit for drug treatment at 10µM for 48h in duplicates. Artemisinin-treated and DMSO-treated cultures were used as positive and negative controls respectively. After 48h, parasitemia was checked through light microscopy. Identical experiments were performed using Dd2 strain to identify cross-active compounds for hit prioritization.

Compounds that showed inhibitory activity were then re-tested at 10, 3, and 1µM for 48h to determine dose-dependent effect. Samples were fixed (0.125% glutaraldehyde), permeabilized (0.1% Triton X100/PBS), and stained (0.25% Hoechst) for flow cytometry. Gating and data analysis were performed using Kaluza (Beckman version 2.1).

### Determination of stage-specific inhibition of parasite growth

Highly synchronized ring stage parasites (5 hpi, 2% parasitemia with 2% hematocrit) were incubated with chosen PRB compounds (at 5µM) that showed >50% inhibition at 3µM, until the trophozoites appeared in DMSO-controls (25 hpi). Parasitemia was determined through flow cytometry with a minimum of 50,000 recorded events. Similar experiments were conducted with trophozoite (25 hpi), as well as schizont stage parasites (35-40 hpi). In parallel, Giemsa-stained samples were inspected for phenotypic consequences upon the drug treatment (20). All experiments were performed a minimum of three times in triplicates to confirm the stage-specific activity.

### Dose-response assay and IC_50_ determination

A serial dilution 96-well plate assay was adopted to determine the IC_50_ of chosen compounds against sorbitol synchronized 3D7 and artemisinin-resistant R539T parasites. The drugs were introduced in 1:2 serial dilutions starting from 10µM to 4.5nM for 48h, in triplicates and parasitemia was scored using flow cytometry. Sigmoidal growth curves were constructed to determine the IC_50_ using non-linear regression analysis using GraphPad Prism (20). Similarly, the Rupture_50_ (concentration at which half the parasite population failed to egress from iRBCs) was determined for the schizont stage inhibitors using the serial dilution method (21).

### Drug combination assay to evaluate synergistic activity

The combinatorial effect of the selected compounds with *artemisinin* and *chloroquine* was investigated using a fixed ratio interaction assay (22). Synchronous ring-stage parasites were treated at appropriate ranges of concentrations such as artemisinin (28nM to 5.6nM), chloroquine, MMV1580496, and MMV1580844 (all between 0.5µM to 2.5µM), MMV1580483 (between 625nM to 125nM), and MMV010014 and MMV1593540 (1µM to 5µM). Drug combinations were prepared for ART-MMV010014, CQ-MMV010014, ART-MMV1593540, CQ-MMV1593540, ART-MMV1580496, CQ-MMV1580496, ART-MMV1580844, CQ-MMV1580844, ART-MMV1580483, CQ-MMV1580483 in fixed ratios (5:0, 4:1, 3:2, 2:3, 1:4, 0:5). This was followed by two-fold serial dilutions of each well and ensuring that the IC_50_ of drug alone (5:0 and 0:5) falls approximately at the mid-way mark. Each well was seeded with parasites at 2% parasitemia and 2% hematocrit to obtain six desired final concentrations for each combination. These experiments, done in triplicates, were allowed to run for 48h. Parasitemia was counted by flow cytometry and analyzed using GraphPad Prism by non-linear regression (sigmoidal dose-response) to yield the IC_50_ for the drug alone and of its combinations. The Fractional Inhibitory Concentration (FIC_50_) for each fixed dose ratio was calculated using the following equation:

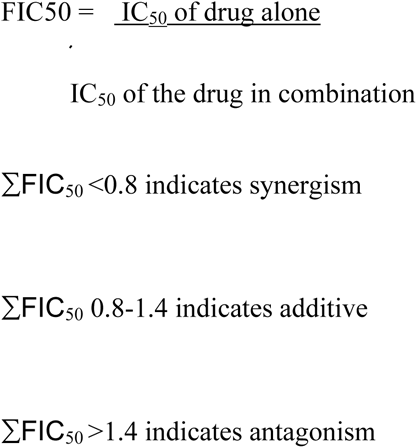

Complete set of data with drug combinations, ratios and overall FIC50 calculation are summarized in **Supplementary Table. 1.**

### Characterization of schizont stage phenotypes using confocal microscopy

The late-stage parasite arrest phenotypes were analyzed by confocal microscopy. To do this, healthy RBCs were pre-stained with Carboxyfluorescein succinimidyl ester (CFSE) fluorescence stain at 5µM, followed by the introduction of MACS-enriched schizonts for infection. The inhibitors were introduced (at 5 µM) around 38 hpi alongside DMSO and E64d as the negative and positive controls respectively. After 15h when rings appeared in DMSO-treated wells, the samples were glutaraldehyde-fixed, permeabilized, and stained with DAPI for confocal microscopy. A Zeiss-LSM980 Airyscan II microscope was used for imaging. High-resolution images were processed using *Zen 3.7* software.

### Measurement of ROS generation in drug-treated parasites

Magnet-purified schizonts (42-44 hpi) were treated with MMV1580483 (1µM), MMV1580173 (1µM), and MMV1634402, MMV1580496, MMV1593540, MMV1580844 (all at 5µM), for 1h, chosen based on their overall inhibitory activity. 1mM tert-butyl hydroperoxide (TBHP) and 0.2% DMSO were used as positive and negative controls, respectively. CellRox green reagent at 5µM (Thermo) was added to the cells for 30 min at 37°C. Cells were co-stained with Hoechst during the last 15 min of incubation in MCM at 37°C in dark. Cells were then washed with PBS three times and processed for flow cytometry. Three independent experiments were conducted with experimental replicates, counting a minimum of 50000 cells for each sample. The P value and significance were estimated using non-parametric ANOVA analysis in GraphPad Prism.

### Recrudescence assay

Ring-stage parasites at 2% parasitemia were treated with MMV001014 at 10µM. After 12h, the cultures were washed and allowed to progress, with untreated parasites serving as controls. The resulting parasitemia at 24, 48, 72, and 96h were quantified using flow cytometry and microscopy. The same procedure (23) was followed for another set of parasites that were treated with the drugs for a duration of 48h which was then tracked for recrudescence over a 30 day period.

### Determination of toxicity of the prioritized compounds

PRB library compounds were originally verified as non-cytotoxic prior to making them available as open resource. Yet, to ensure the selectivity of our hits against *Plasmodium*, an MTT assay was conducted using Vero cells at a single concentration of 20µM, at 15,000 cells/well in triplicates for 24h at 37°C. After the incubation, 10µl of the MTT labelling reagent (Roche 11465007001, 0.5mg/ml) was added to each well and kept for 4h. 100µl solubilization solution was added to each well and kept for 24h for complete solubilization of formazan crystals. Absorbance was measured using a Varioskan Lux Multimode plate reader (Thermo Scientific) at 590 nm (24).

## RESULTS AND DISCUSSION

### Activity determination of PRB library compounds against blood stage *P. falciparum*

Initially, all 400 compounds from PRB were screened against synchronous ring stage 3D7 and Dd2 parasites at a single concentration of 10µM. After 48h, the overall killing efficacy was analyzed through microscopy-based counting from Giemsa-stained smears. From this, sixty compounds inhibiting both strains with more than 90% killing efficacy at 10µM were identified **(Figure. 1)**. All in all, 325 compounds showed limited or no inhibitory potential against either strains, while 15 of them were selective only to any one strain. Only the 60 compounds active against both 3D7 and Dd2 in 3 independent experiments were chosen for further analysis.

**Figure. 1:**
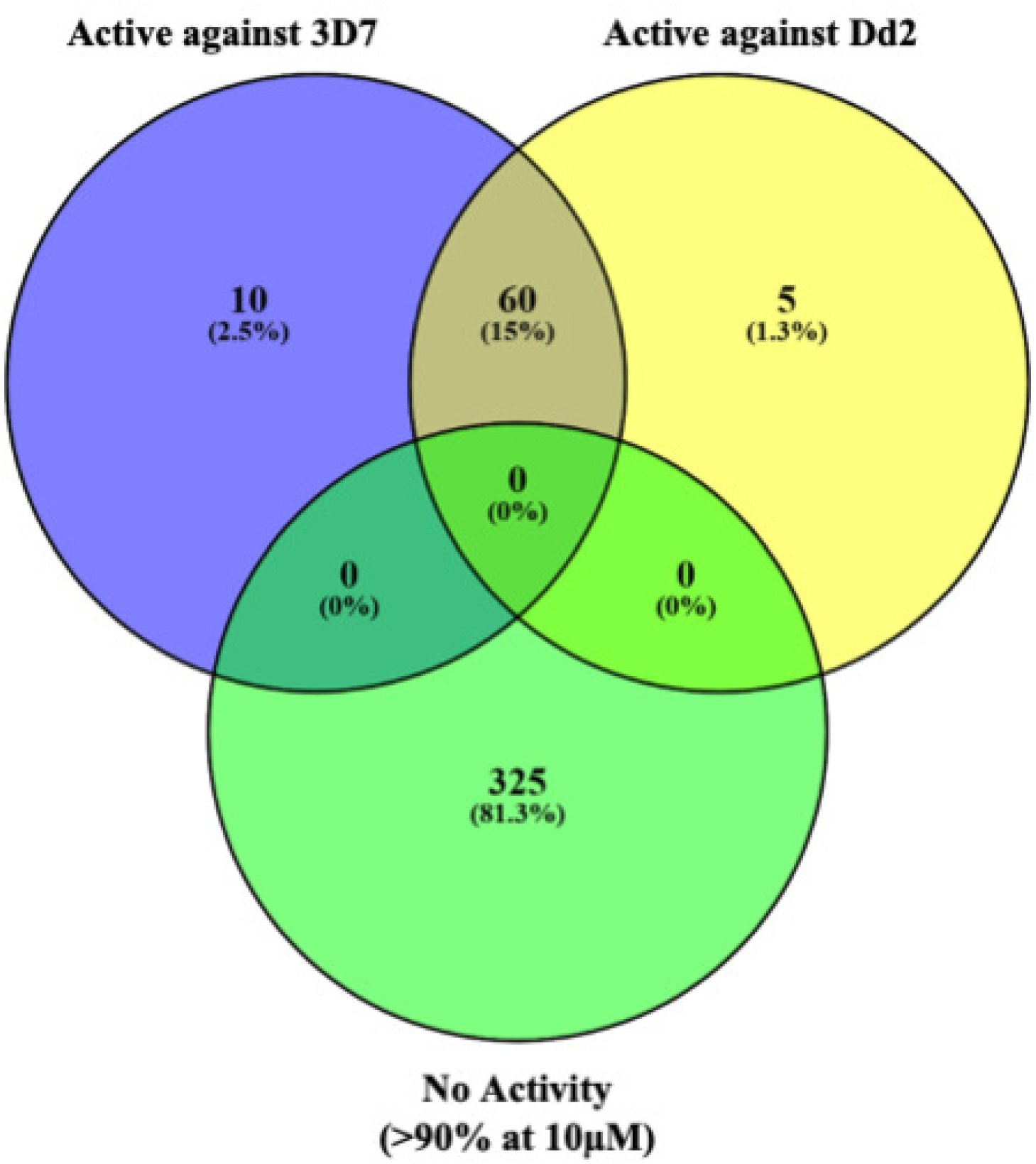
Venn diagram indicating the inhibitory potential of Pandemic response box molecules against blood-stage *Plasmodium falciparum*. Synchronous cultures of 3D7 and Dd2 parasites were separately exposed to the PRB library compounds at a single concentration of 10µM and parasite survival/death was quantified by microscopy. The blue circle represents the drugs active against 3D7 (70), the yellow circle denotes drugs active against Dd2 (65), and the green circle (325) depicts the compounds which showed limited inhibitory activity. 60 compounds were identified to be active against both 3D7 and Dd2 strains.

Next, the compounds were introduced to parasites at 10, 3, and 1µM to estimate the dose dependence. For this, synchronous ring-stage 3D7 cultures were treated for 48 h, followed by parasitemia estimation using flow cytometry. Based on this, 28 compounds active at all concentrations, demonstrating >50% killing efficacy even at 3µM (**Figure. 2A**) were identified. To ensure that these compounds were not cytotoxic, MTT assay was carried out using Vero cells at a single concentration of 20µM (**Figure. 2B**). Overall, the compounds were not cytotoxic in agreement with the information available with the MMV, except for MMV157870, which was omitted from further analysis along with reference compounds *chloroquine* and *tafenoquine,* which were part of the 28 short-listed compounds.

**Figure. 2:**
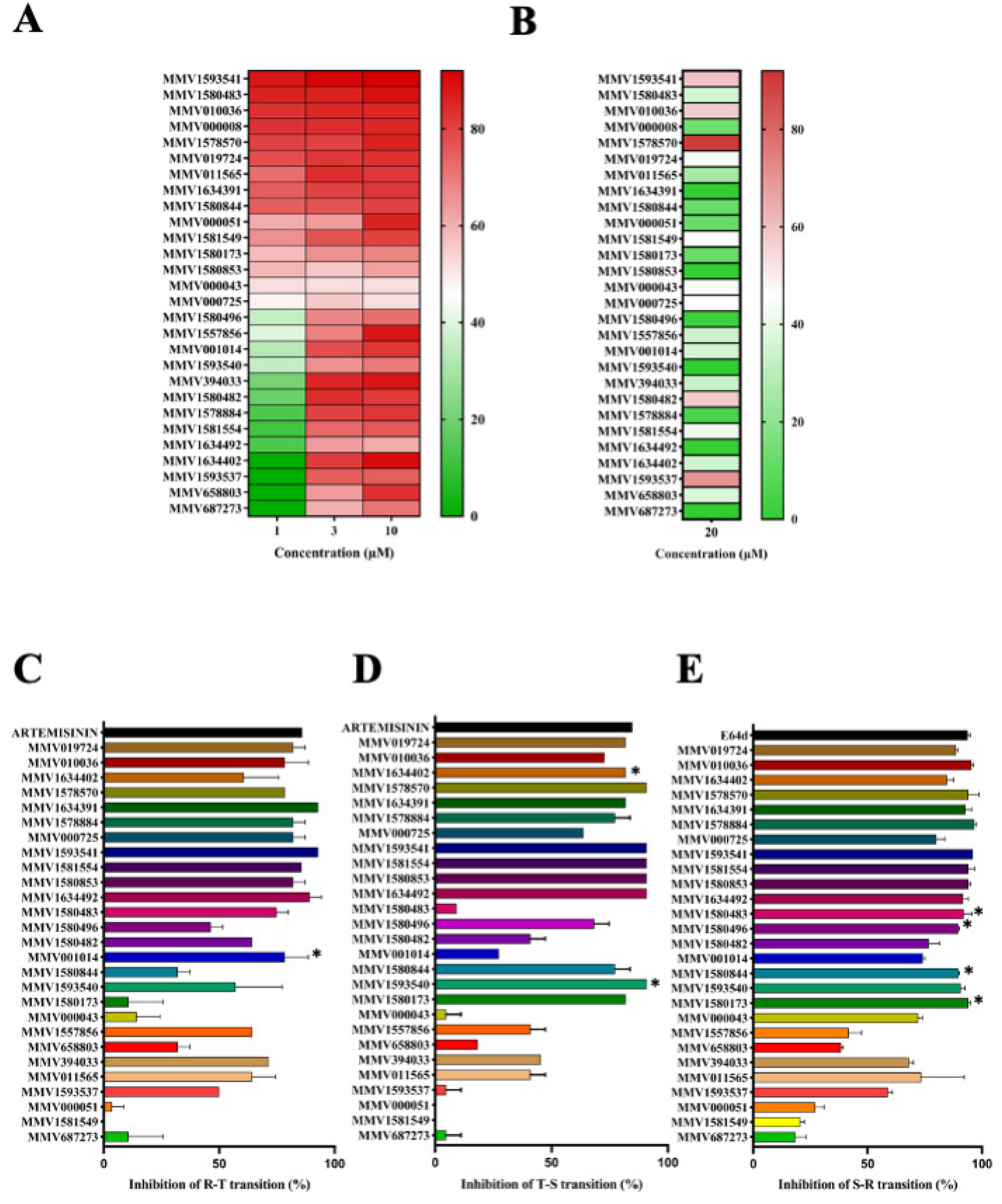
Evaluation of dose-dependent inhibition blood-stage *P. falciparum* by PRB Library compounds. 60 compounds identified from the preliminary screens were re-screened against 3D7 at 10, 3 and 1µM. **(A)** Heat map showing the inhibitory activity of the top 28 compounds; green indicates no inhibition and red indicates highest inhibition. Data is representative of 3 independent experiments performed in triplicates **(B)** Cytotoxicity of the top 28 hit compounds against Vero cells at 20µM determined through MTT assay. The inhibitory potential, expressed as % inhibition relative to non-treated control, of top compounds against parasite transitions; **(C)** ring to trophozoite, **(D)** trophozoite to schizont, and **(E)** schizont to ring. Error bars represent standard deviation.

### Phenotype-based screening to determine stage-specific inhibition of parasite transitions

Growth inhibition assays during stage transitions (such as ring-to-trophozoite, trophozoite-to-schizont, and schizont-to-ring) were carried out, followed by determination of abrogation of development through flow cytometry combined with light microscopy. Ten molecules were found to inhibit all the stages with more than 80% inhibition at 10µM (MMV1634391, MMV010036, MMV019724, MMV1634492, MMV1580853, MMV1581554, MMV1593541, MMV000725, MMV1578884, and MMV1578570), highest potency during the trophozoite stages, with IC_50_ in the nanomolar to low micromolar range **(Supplementary Figure. 1A-C)**. Seven other molecules (MMV001014, MMV1593540, MMV1634402, MMV1580173, MMV1580483, MMV1580496, and MMV1580844) inhibited specific stages (either one of the ring, trophozoite or schizont stage) with more than 90% killing efficacy **(Figure. 2C-E).** Among them, MMV001014 primarily inhibited the ring stages. Two other molecules (MMV1593540 and MMV1634402) abrogated trophozoite stage development, and four molecules (MMV1580844, MMV1580496, MMV1580173 and MMV1580483) halted schizont stage development **(Figure. 2C-E).** A summary of the various stage-specific inhibitors along with their structures and possible mechanisms based on available literature is presented in **Table. 1**.

**Table. 1:**
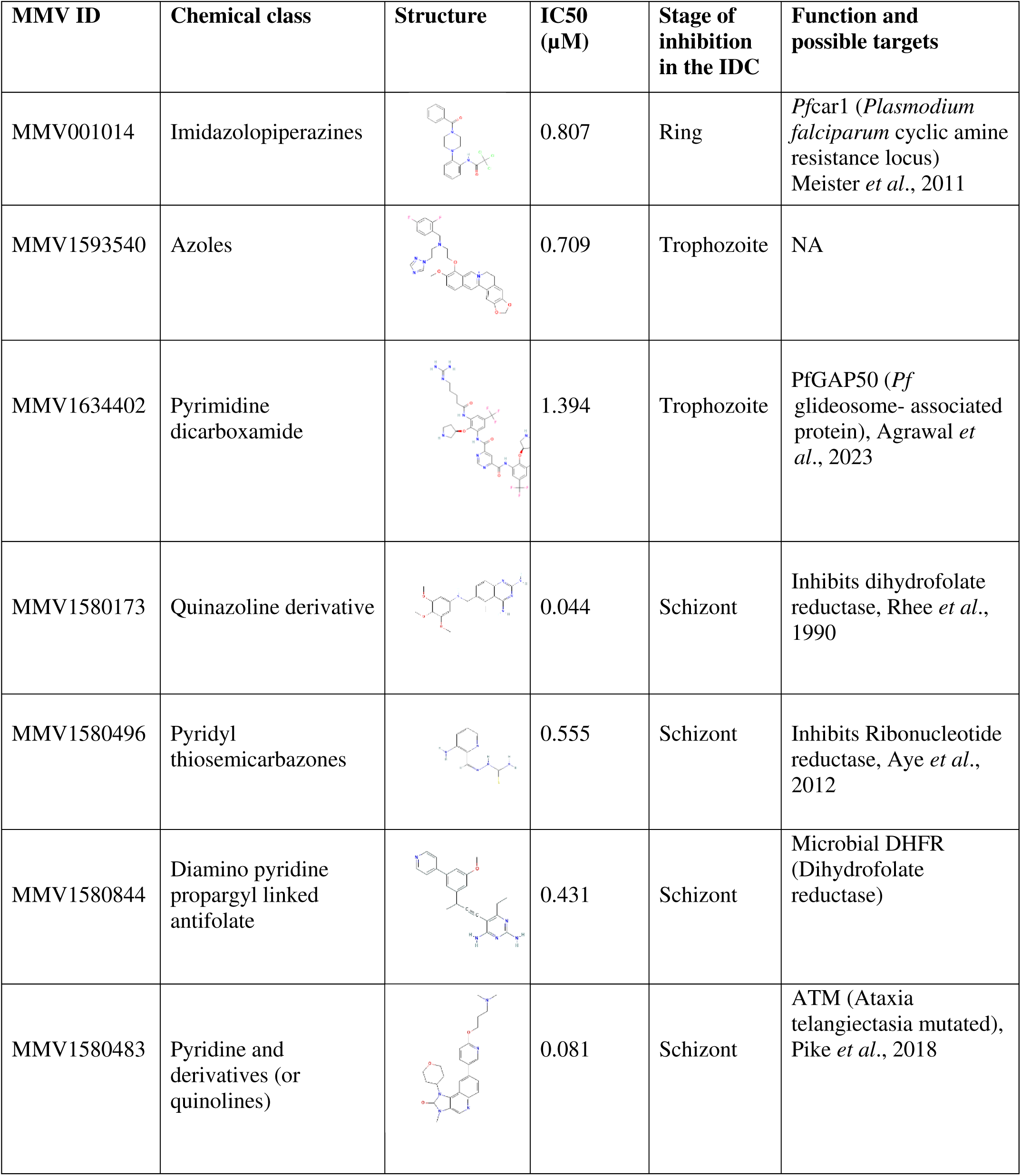
Compiled information of 7 stage specific inhibitors emerged from this screening together with their known mechanisms.

| MMV ID | Chemical class | Structure | IC50 (μM) | Stage of inhibition in the IDC | Function and possible targets |
| --- | --- | --- | --- | --- | --- |
| MMV001014 | Imidazolopiperazines |  | 0.807 | Ring | *Pfcar1* (*Plasmodium falciparum* cyclic amine resistance locus)<br>Meister *et al.*, 2011 |
| MMV1593540 | Azoles |  | 0.709 | Trophozoite | NA |
| MMV1634402 | Pyrimidine dicarboxamide |  | 1.394 | Trophozoite | PfGAP50 (*Pf* glideosome- associated protein), Agrawal *et al.*, 2023 |
| MMV1580173 | Quinazoline derivative |  | 0.044 | Schizont | Inhibits dihydrofolate reductase, Rhee *et al.*, 1990 |
| MMV1580496 | Pyridyl thiosemicarbazones |  | 0.555 | Schizont | Inhibits Ribonucleotide reductase, Aye *et al.*, 2012 |
| MMV1580844 | Diamino pyridine propargyl linked antifolate |  | 0.431 | Schizont | Microbial DHFR (Dihydrofolate reductase) |
| MMV1580483 | Pyridine and derivatives (or quinolines) |  | 0.081 | Schizont | ATM (Ataxia telangiectasia mutated), Pike *et al.*, 2018 |

### Characterization of the ring stage-specific inhibitor MMV001014

In order to further characterize the ring-specific MMV001014, flow cytometry coupled with microscopy was performed. Distinct ring-trophozoite inhibition **(Figure. 3A)** with dead-looking rings were observed through microscopy **(Figure. 3B)**. The estimated IC_50_ for MMV001014 was at sub-micromolar level for 3D7 (0.81µM) while marginally higher for R539T strain (IC_50_ of 1.53µM) **(Figure. 3B).** A washout experiment was conducted to check the reversibility of MMV001014-induced parasite arrest, which showed irreversible action at 6h or 12h exposure followed by washout **(Supplementary Figure. 2A).**

**Figure. 3:**
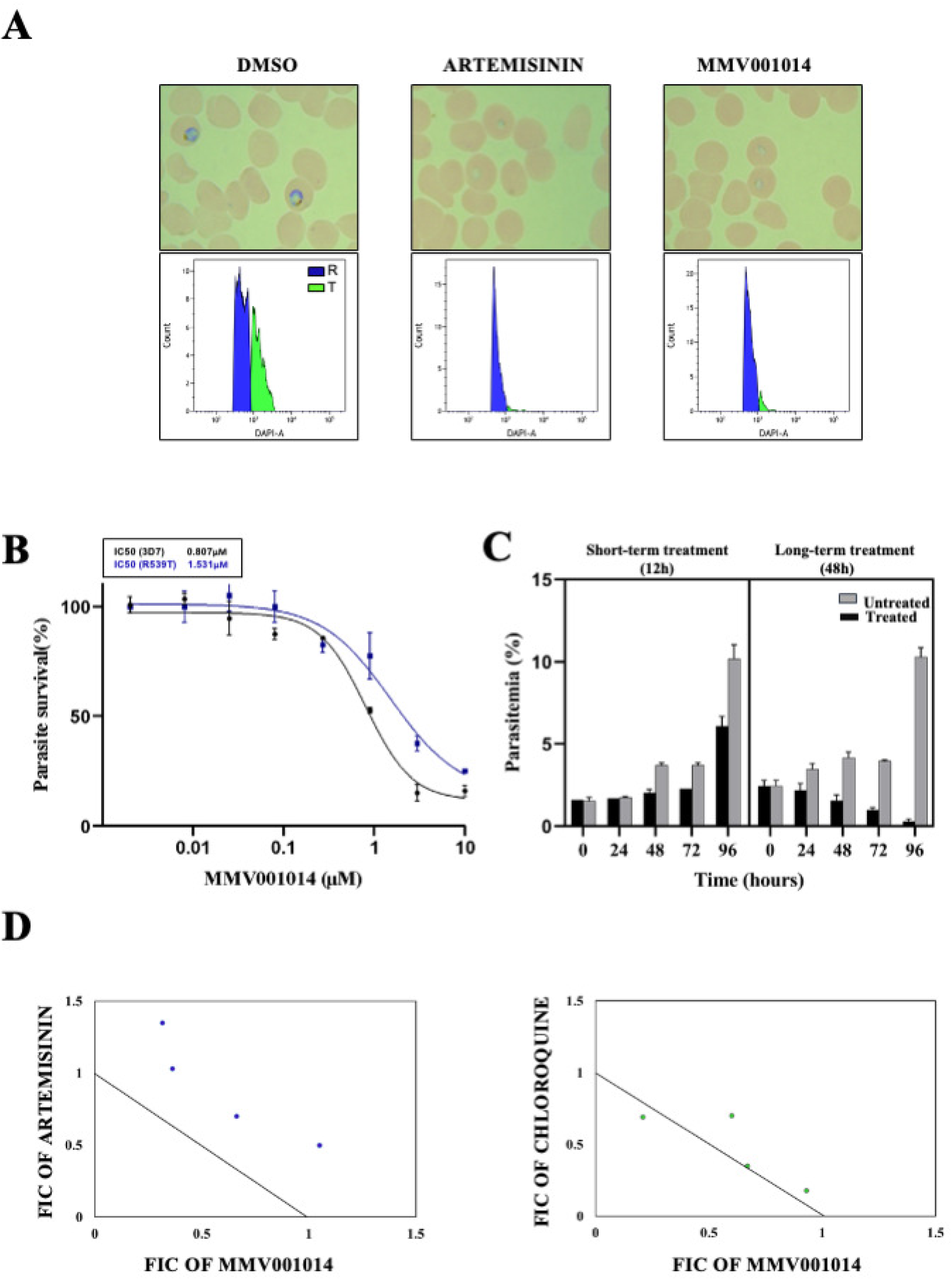
Characterization of the ring-stage specific potency of MMV001014. **(A)** Inhibitory potential of MMV001014 against ring stage parasites was determined by exposing them to the compound at 5µM followed by microscopic analysis of Giemsa-stained smears along with DMSO and artemisinin as controls. Dead rings and complete absence of rings maturing to trophozoites were noted upon treatment with MMV001014, confirmed through microscopy and flow cytometry. In the flow plots, blue colour indicates rings and green colour indicates trophozoites. **(B)** A dose Vs response curve was generated for MMV001014 against 3D7 and R539T parasites through non-linear regression analysis to determine IC_50_ confirming different levels of sensitivity. **(C)** A simplified recrudescence assay was conducted for MMV001014 and the parasite count obtained through flow cytometric analysis (50,000 events) is shown. The left panel indicates drug-treated and untreated samples for short duration (12h), while the right panel shows the drug-treated and untreated samples for a longer duration (48h). In either conditions, no recrudescence was observed. The error bars represent the mean and standard deviation analysed using Graph pad prism. **(D)** Isobolograms summarizing the results of combination treatment of MMV001014 with artemisinin (Left panel) and chloroquine (Right panel) demonstrating antagonistic and additive actions respectively.

To further understand the ability of MMV001014 to impair ring stages permanently and to evaluate possible recrudescence, a single exposure to MMV001014 at 10µM for 12h (followed by removal by washing) was performed. We observed the presence of growth-arrested rings until 48h. In the controls (non-treated), 10% parasitemia was attained within 96h, while only 5-6 percent parasitemia was found in the 12h drug-exposed sample. In another set of parallel samples where prolonged treatment of drugs (for up to 48h) was performed, no healthy parasites were observed. To evaluate the potential of MMV001014 as a partner drug to CQ and ART, a combination assay was adopted (22). This revealed a predominantly antagonistic action of MMV001014 with artemisinin with an FIC value >1.4. In contrast, an additive behaviour (with an FIC close to 1) was recorded for the CQ-MMV001014 combination **(Figure. 3D-E).**

MMV001014 belongs to the imidazolopiperazine family, which forms a distinct scaffold in contrast to the current antimalarials such as aminoquinolines and endoperoxides (25). This class of molecules are broadly active against early blood stage development as well as liver and mosquito stages with some of the related compounds; KAF156 and GNF179 undergoing clinical trials (26, 27). Whole genome analysis showed that resistance to imidazolopiperazines is mediated by mutations in three different genes (*pfcarl*, the cyclic amine resistance transporter, *pfact*, the Acetyl-CoA transporter and *pfugt*, the UDP-galactose transporter), all membrane transporters localized in the endoplasmic reticulum which may inhibit the protein trafficking (28).

MMV001014 showed a predominant ring-stage killing efficacy in our screens, which is the most vulnerable phase of the parasite development, as dead or damaged rings can be easily cleared off from the circulation (29). We observed that MMV001014 being irreversible, which can sustain its potential for a prolonged period if used clinically. Artemisinin being active against ring forms, the antagonistic activity of MMV001014 signifies some level of overlap in their mode of action. Furthermore, the artemisinin-resistant strain (R539T-clinical isolate from Thailand, with mutated *Kelch* gene) showed reduced sensitivity to MMV001014 in alignment with this observation. Chloroquine, on the other hand, mainly affects metabolizing stages such as trophozoites and early schizonts, affecting mainly the metabolic functions. We inferred that MMV001014 may interfere with some of the targets of artemisinin, while there is no interference with chloroquine.

Interestingly, MMV001014 inhibits the mid-late ring stage compared to the artemisinin derivatives (which kill early rings). The recrudescence assay (12h treatment of rings with compound followed by washing) highlighted a clear delay in the recovery of the parasites. At the same time, prolonged drug exposure (48h) resulted in no parasites even after 30 days. The sustained lack of resurgent parasites implies the limited possibility of rapid resistance building if MMV001014 is pursued for clinical applications.

### Growth perturbations induced by trophozoite-specific inhibitors

MMV1593540 and MMV1634402 showed profound inhibitory potential against trophozoite development, further confirmed through microscopic examination of phenotypes **(Figure. 4A)** as well as flow cytometry **(Figure. 4B).** In DMSO-treated controls, schizonts were detected after 15h of treatment while only a few schizonts were observed in MMV1593540 or MMV1634402 treated wells. Although these compounds have been noted for their antimalarial activity (29–31), little about their stage-specific action and mechanisms are known. MMV1593540 is an azole compound active against trophozoites and young schizonts, similar to chloroquine and piperaquine (32). MMV1634402, also known as *brilacidin* is an amphipathic aryl amide foldamer which is currently undergoing clinical trials (33), with *Pf*GAP50 known as the Glideosome-associated protein being its primary target. While *Pf*GAP50 is part of the apical organelles required for host-invasion, prior studies have shown that its transcription is initiated during the ring stage, peaking in the mid-trophozoite stage when compared to the other IMC components such as *Pf*GAP45 and MyoA (34). This is consistent with trophozoite specific inhibitory potential of MMV1634402. It was also interesting to note that neither of these compounds induced compromised digestive vacuole phenotype (35), typical of compounds killing trophozoites. While the inhibition caused by MMV1593540 appeared mostly reversible, MMV1634402 was irreversible indicating a temporally defined interaction with the target **(Supplementary Figure. 2B).**

**Figure. 4:**
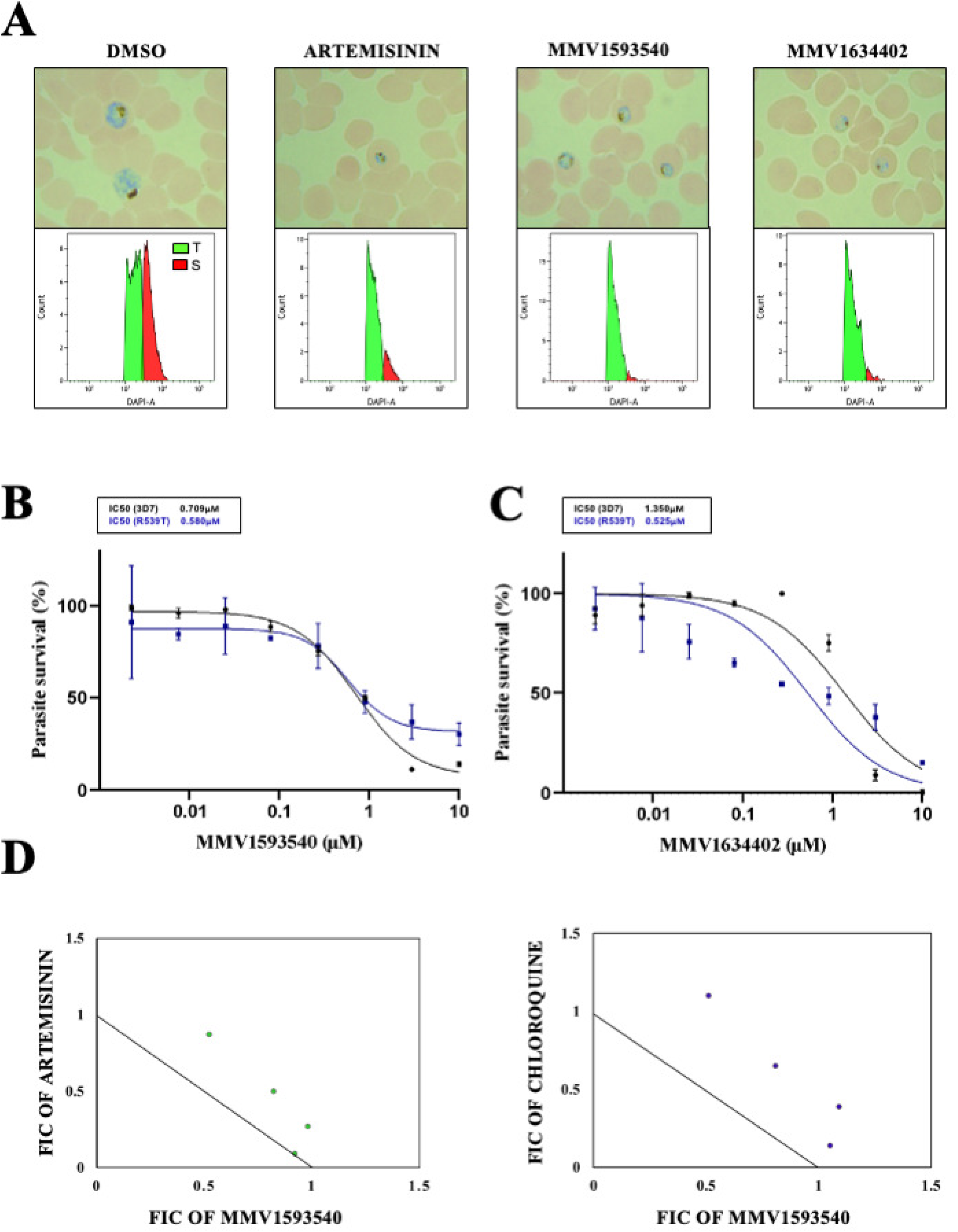
Characterization of the trophozoite-specific inhibitors, MMV1593540 and MMV1634402. **(A)** Microscopic images of parasites upon treatment with MMV1593540 and MMV1634402 (at 5µM) at the trophozoite stage are presented along with controls artemisinin and DMSO-treated controls. Corresponding flow cytometry data is presented as histograms, with green depicting the trophozoites and red indicating the schizonts. The dose vs response curve used for IC_50_ determination against **(B)** 3D7 and **(C)** R539T strains obtained through non-linear regression analysis using graphpad prism are represented. **(D)** The combinatorial effect of MMV1593540 and MMV1634402 with artemisinin and chloroquine are represented as isobolograms respectively. MMV159350 showed additive action with artemisinin while antagonistic action with chloroquine.

MMV1593540 showed an IC_50_ of 0.7µM against 3D7 and 0.58µM against R539T (**Figure. 4B**). In contrast, corresponding numbers for MMV1634402 (*Brilacidin*) were 1.35µM and 0.53µM respectively **(Figure. 4C)**. As such, it appeared that the trophozoite inhibitors have superior efficacy against ART-resistant parasites which is noteworthy. Significantly high IC_50_ of MMV1634402 was perhaps due to the unique host defence protein mimetic structure limiting its permeability across the membranes surrounding the parasites (30). More research is needed however to explain this observation. In the combination studies, MMV1593540 demonstrated partially additive action which indicate a lack of overlap with ART while antagonistic action with CQ **(Figure. 4D).** These compounds also showed significant ROS generation **(Supplementary Figure. 3)** within one hour of treatment, which may be one of the modes of action or a secondary mechanism of the drugs inducing cell death.

### Characterization of schizonticidal compounds from PRB

We identified 4 compounds namely MMV1580173, MMV1580844, MMV1580496 and MMV1580483 inhibiting schizont maturation (or progression to subsequent ring stages), with different degrees of reversibility **(Supplementary Figure. 2C).** Late-stage schizonts were visible in drug-treated samples for several hours after rings were observed in the DMSO-treated control **(Figure. 5A).** All of them, including the Dihydrofolate reductase *(DHFR)* inhibitor Trimetextrate (MMV1580173) showed excellent schizonticidal activities with IC_50_ in the nanomolar range **(Figure. 5B-E).** MMV1580483 (AZD-0156) was the most potent schizonticidal compound with an estimated IC_50_ of 40nM and 79nM against 3D7 and R539T respectively. Furthermore, all of these inhibitors displayed predominantly antagonistic action with artemisinin and chloroquine except MMV1580844 which showed additive action with artemisinin **(Figure. 5F-H),** indicating vastly different modes of action.

**Figure. 5:**
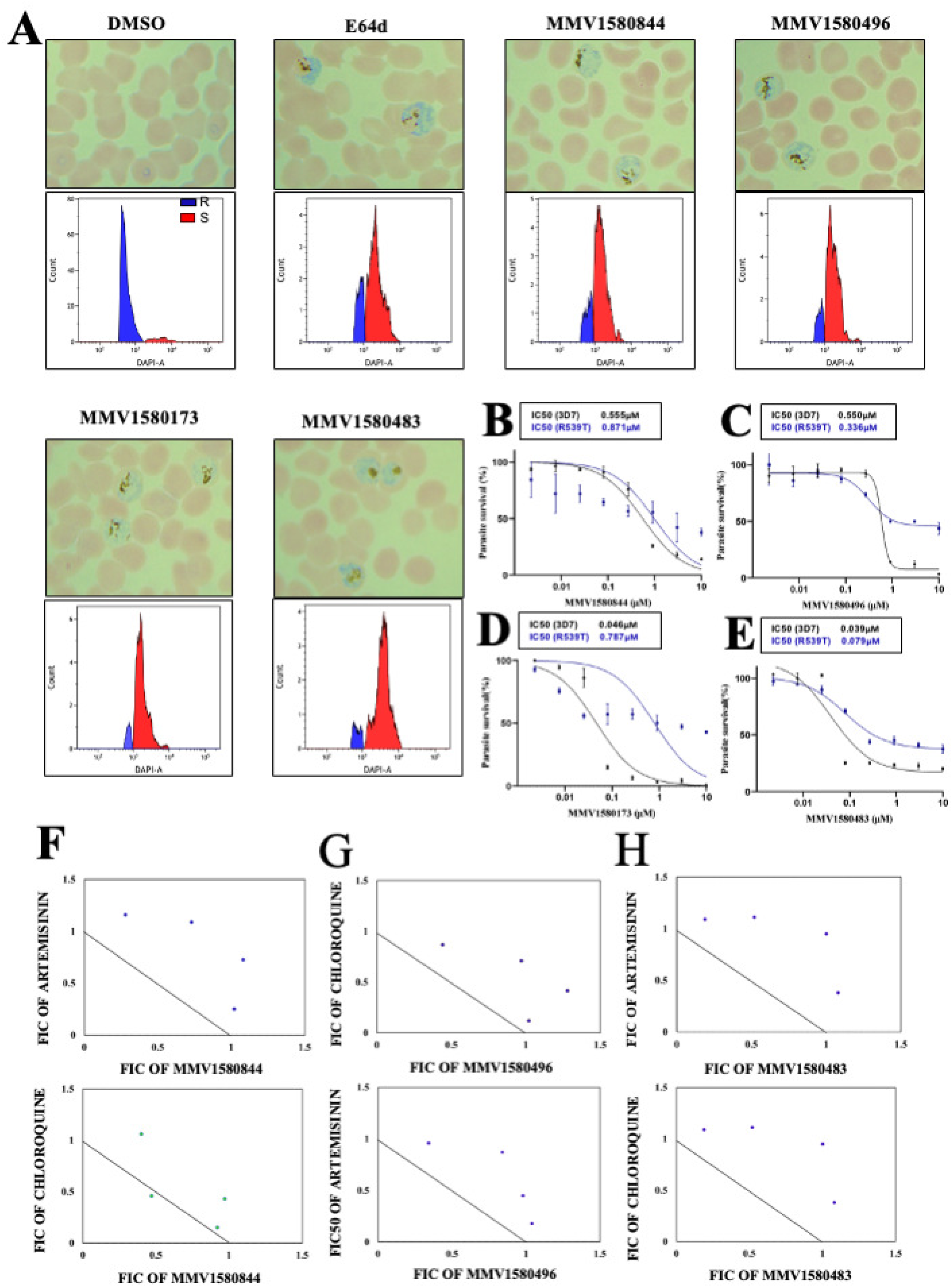
Characterization of schizont arrest induced by PRB Box compounds. **(A)** Microscopic images of growth-arrested schizonts induced by MMV1580173 (at 1µM), MMV1580844 (at 5µM), MMV1580496 (at 5µM), and MMV1580483 (at 1µM), along with the corresponding flow cytometry patterns (Blue: ring stage parasites, Red: schizont stage parasites). E64d and DMSO were used as positive and negative control respectively. **(B-E)** The drug sensitivity of the schizont stage inhibitors against 3D7 and R539T strains are indicated along with their IC_50_ values. The combinatorial effect of these compounds with artemisinin and chloroquine are represented in figures 5F-H. MMV1580844 showed antagonistic action with chloroquine while additive action with artemisinin, while MMV1580496 and MMV1580483 show antagonistic action with chloroquine and artemisinin.

MMV1580173 belong to the class of quinazoline derivatives, is an already known antimalarial targeting dihydrofolate reductase. The core pyramidine ring structure of the compound makes it active against DHFR killing early schizonts. It needs to be however noted that the substituent group (methyl piperazine) makes it more specific against human DHFR compared to prokrayotic DHFR (36). In contrast MMV1580844 is a diaminopyridine propargyl linked antifolate, shown to possess selective inhibition against prokaryotic DHFR compared to human DHFR, with established antimicrobial activity (37, 38). MMV1580496, commonly known as Triapene, belonging to the class of thiosemicarbazones, is an iron-chelator and thought to inhibit the *ribonucleotide reductase* through the production of ROS (39). The ribonucleotide reductase catalyzes the oxidation of nucleotide diphosphates (NDPs) into deoxynucleotide diphosphates (dNDPs), in pyrimidine and purine pathways. RNR subunits R1 and R2 reach their peak transcriptome level at 35 hpi in the schizont stage. Triapene has shown killing activity against *Plasmodium* and *Toxoplasma spp* (39, 40), although its target and mode of action remains unclear. MMV1580483, commonly known as AZD-0156, belonging to the class of quinolines, is an ATM kinase inhibitor and a chemo-sensitizing agent against cancer cells (41).

To differentiate the ability of these compounds to arrest late stage schizont development and/or parasite release, we performed time-resolved egress invasion assays from which Rupture_50_ (drug concentration at which half of the schizonts failed to exit the iRBCs) was calculated (20). This analysis indicated that the compounds showed egress/invasion inhibitory activity at a concentration more or less similar to their overall IC_50_ **(Figure. 6A).** Next, confocal microscopy was performed on schizont stage iRBCs pre-stained with CFSE (before infection) along with DMSO and E-64d (egress inhibitor) treated parasites. Parasites inside the iRBCs were stained with Hoechst, which provided us with early insights into parasitophorous vacuole and iRBC membrane integrity **(Figure. 6B).** Interestingly, parasites treated with MMV1580173 were distributed throughout the iRBC with dispersed and fuzzy staining for DNA indicative of overall merozoite disintegration. In contrast, merozoites were intact when schizonts were treated with any of MMV1580844, MMV1580483, or MMV1580496. However, the CFSE staining was mostly absent in these phenotypes, indicative of a compromised iRBC membrane. High-resolution z-stack images of the iRBC obtained through confocal microscopy confirmed that the compounds were killing the parasites at the schizont stage, with different degrees of iRBC compromise.

**Figure. 6:**
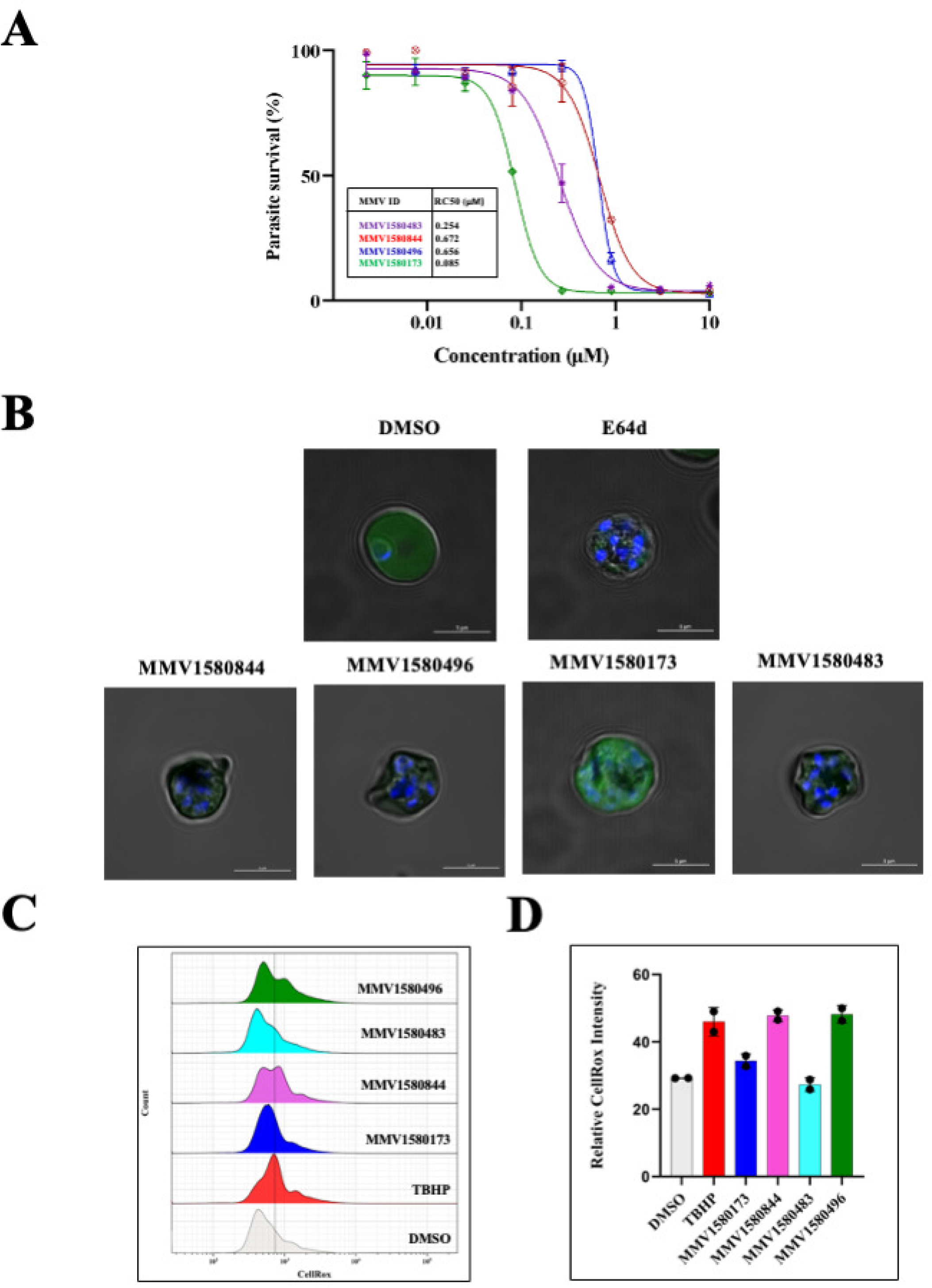
Phenotypic characterization of the schizonticidal inhibitors. **(6A)** Represents the dose-response curve showing the inhibition of parasite egress arrest indicated as Rupture_50_ (concentration required for 50% rupture of infected cells), for the schizonticidal agents MMV1580173, MMV1580844, MMV1580496, and MMV1580483. **(6B)** Z stack confocal microscopic images of the schizont-infected RBCs treated with compounds confirming the schizont stage inhibition and resulting morphology. **(6C)** ROS generation within one-hour of drug treatment is depicted as flow plots and bar diagrams with relative CellRox fluorescence intensity. TBHP was used as the positive control while DMSO was the negative control. All of the schizonticidal inhibitors led to significant ROS generation with MMV1580483 and MMV1580173 showing the least effect.

Next, we estimated the extent of free radical generation in the inhibitor-treated schizonts to differentiate egress inhibition Vs a general schizont death. For this, ROS generated by the stage-specific inhibitors 1h post-treatment was analyzed using a fluorogenic dye CellRox which detects superoxides and hydroxyl radicals **(Figure. 6D).** We noticed that MMV1580173 and MMV1580483 generated relatively low ROS-induced damage similar to DMSO-treated controls while the effect was consistently more significant for all the other schizonticidal compounds **(Figure. 6E).** For the compounds showing immediate ROS generation, it can be inferred that their mechanism likely involves oxidative stress, which could either be a mechanism or consequence of an unknown cellular response. At the same time, minimal ROS production indicates that these drugs might operate through target-specific mechanisms, which need targeted and molecule-specific approaches for investigation in future research.

Taken together, our results on the newly identified inhibitors from PRB library and stage-specific phenotypes are indicative of promising drug-like small molecules with unique modes of action which could form the basis of downstream mechanistic understanding for novel antimalarial development.

## SUMMARY

We present a comprehensive dataset on the stage-specific and phenotypic activities of small molecules belonging to the PRB library against blood stage *Plasmodium falciparum* development. Through this analysis, we prioritized candidate molecules that could be exploited for re-purposing to aid new combinations to combat Malaria. Furthermore, life-stage resolved window of activity and early insights into the mode of action derived through this phenotypic documentation will form the foundation to design experimental protocols to identify novel targets against which new chemotherapeutic avenues could be explored.

## Supplementary Figure Legends

**Supplementary Table. 1:** Complete set of information from the various drug combination assays (combinations, ratios and overall FIC50 calculation are summarized in **Supplementary Table. 1.**

**Supplementary Figure. 1: Antimalarial activity of the compounds inhibiting all intra-erythrocytic forms of *Plasmodium*. (A)** Dose response curves of the compounds inhibiting all stages of parasite development is presented. Parasite survival is indicated in the Y axis against drug concentration on the X axis, plotted based on non-linear regression analysis using Graphpad prism. **(B)** The figure represents the 2D chemical structures of the active compounds identified which were retrieved from PubChem.

**Supplementary Figure. 2: Reversibility profiles of the stage specific inhibitors emerged from this screen. (A)** Parasitemia calculated following 3h, 6h and continued exposure for up to 15h to MMV001014 (at 5µM) indicating mostly irreversible inhibition of parasite development. **(B)** Parasitemia calculated following 3h, 6h and continued exposure for up to 15h the trophozoite stage inhibitors MMV1593540 and MMV1634402 (at 5µM) are presented. **(C)** Parasitemia calculated following 3h, 6h and continued exposure for up to 15h to the schizont stage inhibitors MMV1580173 (at 1µM), MMV1580844 (at 5µM), MMV1580496 (at 5µM), and MMV1580483 (at 1µM) are presented.

**Supplementary Figure. 3: Analysis of ROS generation by Trophozoite inhibitors. (A-B)** Figure represents the ROS levels within one hour treatment of trophozoite inhibitors (MMV1593540 and MMV1634402, both (at 5µM)) measured as the relative Cellrox fluorescence intensity using flow cytometry. TBHP was used as the positive controls and DMSO as the negative control.

**Supplementary Dataset: Complete information from the screening of MMV pandemic Library compounds derived in this study.**

## Author Contributions

**ATP:** Involved in study design, performed experiments, analysed the data and assisted with manuscript preparation; **DKM:** involved in the data analysis and preparation of manuscript; **RC:** Designed the study, supervised research, acquired funding and wrote the manuscript.

## Statement of Conflicts of Interest

Authors have no conflicts to declare.

## Acknowledgements

The authors acknowledge the infrastructure and intramural research support through the BRIC Rajiv Gandhi Centre for Biotechnology and Professor Chandrabhas Narayana (Director, RGCB). They also acknowledge the assistance in flow cytometry experiments and analysis by Ms. Surabhi and Mr. Thilak. The authors thank Mr. Anuroop for helping with confocal microscopy and image analysis/interpretation. They acknowledge Dr. Shailja Singh (JNU, New Delhi), Dr. Souvik Bhattacharjee (JNU, New Delhi), Dr Kailash Pandey (ICMR-NIMR, New Delhi) and Dr. Dhanasekaran Shanmugam (CSIR-NCL, Pune) for gifting various parasite strains. Support in performing the cytotoxicity assays (Dr. Sara Jones and Ms. Remya Raveendran, RGCB) is gratefully acknowledged.

